# Polarized *Entamoeba:* A model for stable bleb driven motility

**DOI:** 10.1101/683813

**Authors:** Deepak Krishnan, Sudip Kumar Ghosh

## Abstract

Protozoan parasites Entamoeba histolytica and Entamoeba invadens formed a polarized phenotype, an elongated shape with a single leading edge and a trailing edge when treated with pentoxifylline. The leading edge of the polarized morphology was a spherical protrusion devoid of F-actin but with occasional F-actin scars, indicating the presence of bleb. The polarized form was stable bleb driven since the blebbing was limited to the leading edge. Pentoxifylline induced chemokinesis in Entamoeba as it switched the motility pattern from slow and random to fast and directionally persistent. Pentoxifylline speeded up the cell aggregation in E. invadens during growth and encystation due to enhanced chemotaxis of the polarized form. The transformation of non-polarized adherent trophozoites to nonadherent stable bleb driven form occurred via lamellipodial and bleb driven adherent intermediate phenotypes. The nonadherent polarized phenotype was highly motile under confinement and moved by rearward plasma membrane flow. In contrast to pentoxifylline, adenosine, the adenosine receptor agonist, stimulated the formation of multiple protrusions leading to random motility. Thus pentoxifylline might prevent lateral protrusions by inhibiting adenosine receptor, producing the monopodial polarized morphology.

**Summary statement:** Pentoxifylline, the adenosine receptor antagonist induced a stable bleb driven polarized morphology in *Entamoeba* characterized by fast, directionally persistent and highly chemotactic motility.

## INTRODUCTION

Cell migration is an important event in embryonic development, wound healing, immune response, and cancer progression. The first step in cell migration is the polarization during which the cells break symmetry to form a leading edge and a trailing edge. Depending on the leading edge, motility can be of two types, lamellipodial motility driven by actin polymerization and pressure driven blebbing. Cells like keratocytes and fibroblasts use lamellipodia at the leading edge while primordial germ cells and amoeboid tumor cells use blebbing motility (Blaser *et al.*, 2006; Paňková *et al*., 2010). Bleb driven cell migration have been reported in many biological events like embryonic development, *Dictyostelium* chemotaxis, and cancer metastasis (Fackler and Grosse, 2008) but unlike lamellipodial motility, its mechanics is not well understood.

Enteric parasite *Entamoeba histolytica* is a highly motile organism, and it has been reported to use only the bleb driven motility both *in vitro* and *in vivo* and thus can be considered as an important model to study blebbing (Maugis *et al*., 2010). As shown previously in progenitor cells (Diz-Muñoz *et al*., 2010), blebbing reduced directional persistence in *E. histolytica*. Here we report that both human pathogen *E. histolytica* and its reptilian counterpart and encystation model *Entamoeba invadens* readily formed a fast moving, elongated morphology when treated with a millimolar concentration of methylxanthines like pentoxifylline or caffeine. This morphology showed a single leading edge with stable bleb, making it directionally persistent.

## RESULTS AND DISCUSSION

### Methylxanthines induced stable bleb driven polarized morphology in *Entamoeba*

*Entamoeba histolytica* (*Eh*) and *Entamoeba invadens* (*Ei*) trophozoites are unpolarized *in vitro* growth conditions which were observed from the nearly circular shape (Fig.1A, control) and the high circularity value of 0.87 (Fig.1B, control), a measure of polarity (Lomakin *et al*., 2018). When exposed to a millimolar concentration (0.5-10 mM) of pentoxifylline (Ptx), both the *Entamoeba* species turned into a highly elongated morphology with a single leading edge and a trailing edge called uroid (Fig.1A Ptx; Movie 1) and reduced circularity (Fig.1B Ptx). Ptx induced polarized morphology was characterized by the aspect ratio (ratio of the major axis to the minor axis). Both *Entamoeba* species showed an average aspect ratio of 1.3 in growth medium (Fig.1C Control), and pentoxifylline treatment increased the aspect ratio to 2.27 in Eh and 3.2 in case of *Ei* (Fig.1C Ptx). Phase contrast micrograph indicates that the cytoplasm of these elongated cells is separated into a fluid endoplasm and gelled ectoplasm. At the leading edge, the endoplasm gelled to form stationary ectoplasm and, at the trailing edge, the ectoplasm was again converted to fluid endoplasm, which can be clearly perceived from the movement of intracellular particles (Fig.1D; Movie 2).

**Fig. 1.**
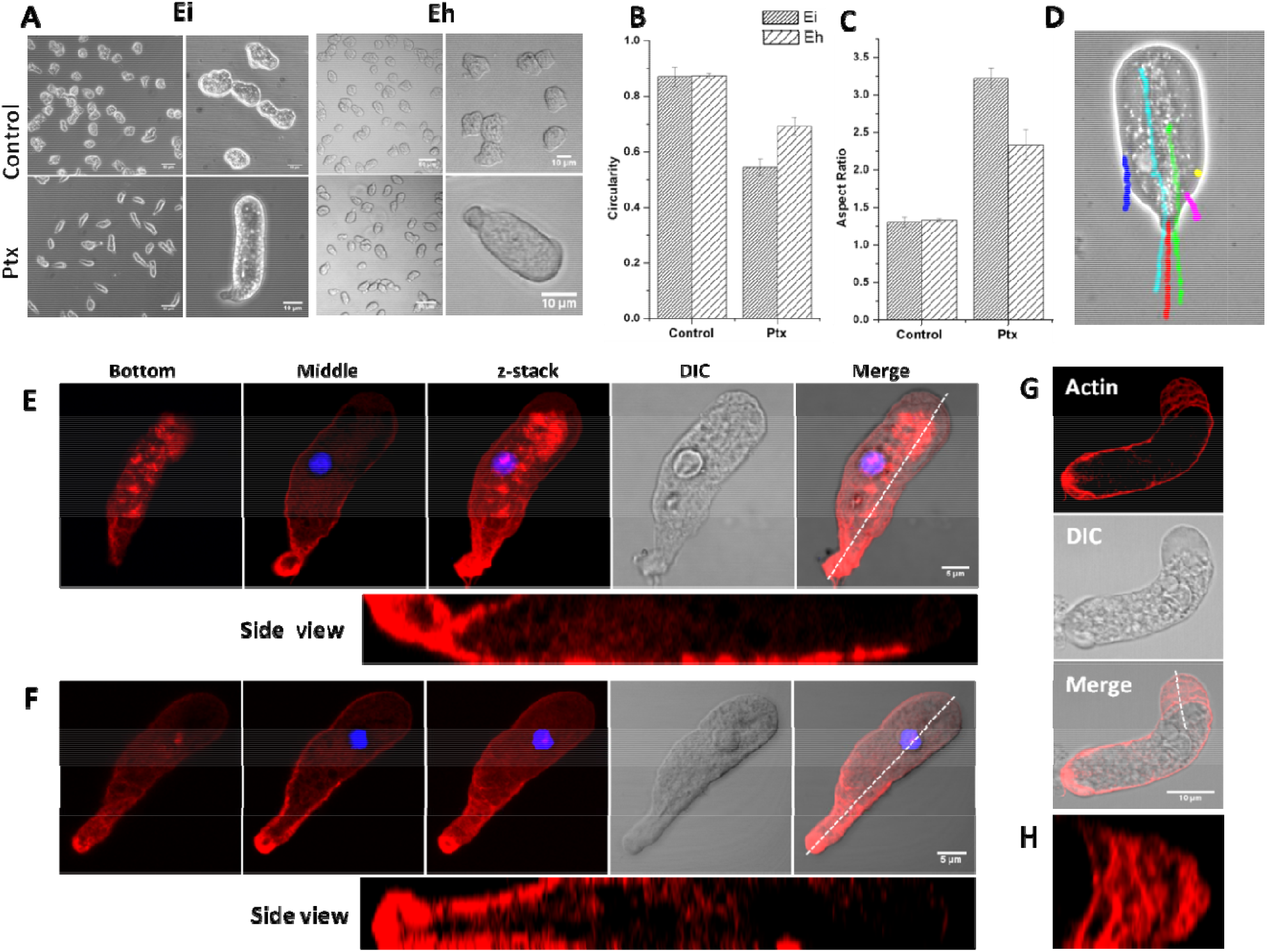
Pentoxifylline (Ptx) induced stable bleb driven polarized morphology in *Entamoeba*. **(A)** Normal and polarized morphology of *E. invadens* (Ei) and *E. histolytica* (Eh). **(B)** Circularity and **(C)** aspect ratio before and after Ptx treatment. **(D)** The cytoplasm of polarized form was divided into a fluid endoplasm and a gelled ectoplasm as shown by the speed of particles (coloured tracks) compared to the cell (red). **(E, F)** In the polarized form, actin was found mainly in the uroid and subcortical regions. Actin fluorescence intensity decreased towards the leading edge, and it was mostly devoid of F-actin. The early polarized form showed actin-based adhesion structures on the ventral side of the cell (E), but they disappeared afterward (F). **(G)** F-actin scars observed at the leading edge of cells changing direction. **(H)** Cross-section of the leading edge. Scale bars: 10 μm.

Since cytoplasmic microtubules are absent in *Entamoeba*, the actin cytoskeleton alone is responsible for their morphology and motility (Vayssié *et al*., 2004). In the unpolarized trophozoites, F-actin was found in the structures like phagocytic invaginations, stress fibers, and podosomes (Fig.S1) (Manich *et al*., 2018). Whereas in the polarized form, most of the F-actin was localized at the uroid and the subcortical region, and its quantity was found to gradually decrease from uroid to leading edge (Fig.1 E, F). Actin-based adhesion structures were initially observed on the ventral side of the cell as they attained polarized form (Fig.1E) but disappeared with time (Fig.1F). Unlike the actin polymerization-driven lamellipodial migration (Fig.S2), the single spherical hyaline protrusion at the leading edge contained the lowest amount of F-actin compared to other regions of the cell. This indicates that the motility of the polarized form was driven by bleb, stabilized at the leading edge. Similar motility was previously reported in zebrafish progenitor cells and cancer cells under confinement and low adhesion (Liu *et al*., 2015; Ruprecht *et al*., 2015). *Eh* used bleb-driven motility, but the blebs were produced all over the cell, causing random motility (Maugis *et al*., 2010). The life cycle of a bleb involves depolymerization of the actin cytoskeleton, causing the intracellular pressure to push the membrane forward, followed by the actin cortex reformation under the plasma membrane. Actin scars from such cyclic generation and healing of actin cortex were observed in many polarized cells, especially those performing a turn (Fig.1 G, H). Thus a single spherical leading edge devoid of F-actin and the occasional presence of F-actin scars also indicate that the motility of Ptx induced polarized cells are stable-bleb driven.

### Motility characteristics of the amoeboid form

The movement of unpolarized and polarized forms of *Entamoeba* was characterized by measuring the velocity and the directness (ratio of displacement to distance, a measure of directional persistence) of the migration (Zengel *et al*., 2011). The velocity of both the *Entamoeba* species in a fresh growth medium was ~0.1 μm/s, and in 12-hour old growth medium, it increased to ~0.3 μm/s (Fig.2A). This change in velocity could be because of the chemokinetic effect of chemicals secreted by the trophozoites (Zaki *et al*., 2006). In both conditions, the cell movement was random as observed from the low directness values (Fig.2B). In the case of Ptx induced polarized form, velocity increased to nearly 1 μm/s (Fig.2A, Ptx). Most importantly, the movement was highly directional with a directness value of 0.7 (Fig.2B, Ptx). These changes in the motility pattern were easily observed from the migrational tracks (Fig.2C; Movies 3, 4).

**Fig. 2.**
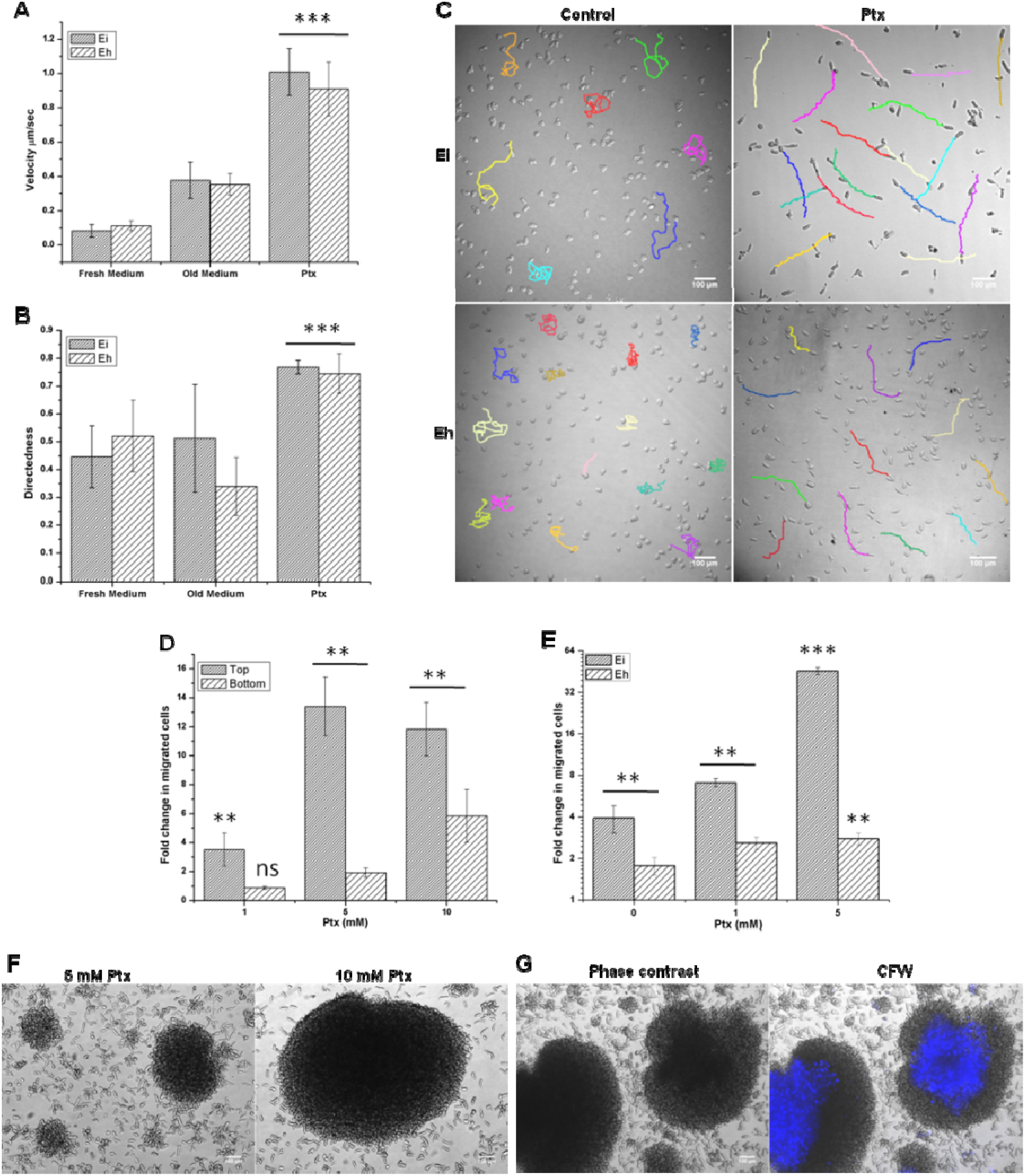
Motility of pentoxifylline (Ptx) induced polarized morphology. **(A)** Velocity and **(B)** Directness *E. histolytica* and *E. invadens* trophozoites in the fresh growth medium, in old medium and after Ptx treatment. **(C)** The tracks of randomly chosen *Eh* and *Ei* trophozoites before, and after Ptx treatment. **(D)** Transwell migration assay showing chemokinetic nature of Ptx. Ptx dose-dependently increased migration but at 10mM Ptx cell aggregation in upper well reduced migration. **(E)** Ptx increased the chemotaxis of serum-starved cells towards serum-containing medium. **(G)** Ptx dose-dependently increased cell aggregation in growth media. **(H)** Encystation medium and Ptx together formed cell aggregates at low cell density and cyst formation inside such aggregates shown by CFW staining (blue). Scale bar: 100 μm.

Transwell migration assay was performed to find out whether Ptx induced motility is due to chemokinesis or chemotaxis. Ptx was added at different concentrations to the bottom chamber to test chemotaxis and to the top chamber to stimulate chemokinesis. Cell migration increased dose-dependently in both the cases, but the migration was much higher when Ptx was added to the upper chamber (10 mM Ptx reduced migration due to extensive cell aggregation in the upper chamber as discussed below) indicating that the effect was due to chemokinesis (Fig. 2D). The most important aspect of Ptx induced chemokinesis was the increased directional persistence due to cell polarization. Factors that increase morphological polarity and directional persistence have been shown to promote chemotaxis (Bosgraaf *et al*., 2005; Pankov *et al*., 2005). Also, blebbing cells were reported to be strongly chemotactic (Zatulovskiy *et al*., 2014). To find the chemotactic potential of polarized *Entamoeba* cells, Transwell migration of serum-starved *Entamoeba* trophozoites towards serum-containing medium was studied. Ptx increased chemotaxis in both *Entamoeba* (Fig.2E), but it was more significant in *Ei*.

*Ei* cysts are formed only within the galactose ligand-mediated cell aggregates in encystation media (Turner and Eichinger, 2007). The Ptx induced polarized morphology was also capable of forming similar galactose ligand-mediated cell aggregates in the growth medium (Fig.2F; Movie 5). Time-lapse video microscopy showed that during *in vitro* encystation, *Ei* formed the polarized morphology in response to encystation media (Movie 6) and these polarized cells moved towards the preexisting aggregates and formed larger aggregates (Movie 7). Therefore, the cell aggregation during encystation might be due to chemotaxis of polarized cells towards the small cells clusters, acting as aggregation centers. Addition of Ptx to encystation medium increased the cell aggregation even at low cell density (10^4^ cells/ml against 10^5^ cells/ml required for *in vitro* encystation), and cysts were also formed in such aggregates (Fig.2G). Ptx also can induce cell aggregation in *Eh*, but these aggregates were short-lived. These differences in the chemotactic and aggregation potential may be two important reasons why the *Ei* aggregated and encysted *in vitro* in response to nutrient and osmotic stress but not the *Eh*.

### Temporal progression of morphological and motility changes during cell polarization

The analysis of morphological changes in *Ei* by actin staining showed that after treatment with 1mM Ptx, phagocytic invaginations on the cell surface (Fig.3A a, a’) started disappearing and the cells started producing lamellipodia (Fig.3A b, b’, c, c’; Fig.S3A; Movie 8). By 4 hours, the lamellipodia became more and more prominent until a polarized form with lamellipodia at the leading edge was formed (Fig.3A d, d’). By the 6^th^ hour, blebs started appearing at the leading edge (Fig. 3A e, e’) and the cell became elongated (Fig. 3A f, f’). Actin-based adhesion structures, similar to the actin feet of *Dictyostelium* (Fukui and Inoue, 1997; Uchida and Yumura, 2004) were found on the ventral side of these initial bleb driven cells (Fig.3A g, g’; S3B). The adhesion structures disappeared by 8-12 hours, giving rise to adhesion independent, stable bleb driven morphology (Fig.3A h, h’). These morphological changes depended on the Ptx concentration, which can be seen in the increment in cell velocity with time (Fig. 3B). Live cell video microscopy showed that the lack of adhesion structures caused the cells to slip while trying to move, which was seen as the reduction in velocity in later hours for higher Ptx concentration (Fig.3B, 10mM, Movie 9). The morphological transition in *Eh* required higher Ptx concentration (5-10 mM) than that of *Ei* (0.5-1 mM). In *Eh* the lamellipodial form was not seen, and the adherent cells (Fig.3C a, a’) produced multiple blebs (Fig.3C b, b’) in response to Ptx and directly formed the stable bleb driven morphology (Fig.3C c, c’).

**Fig. 3.**
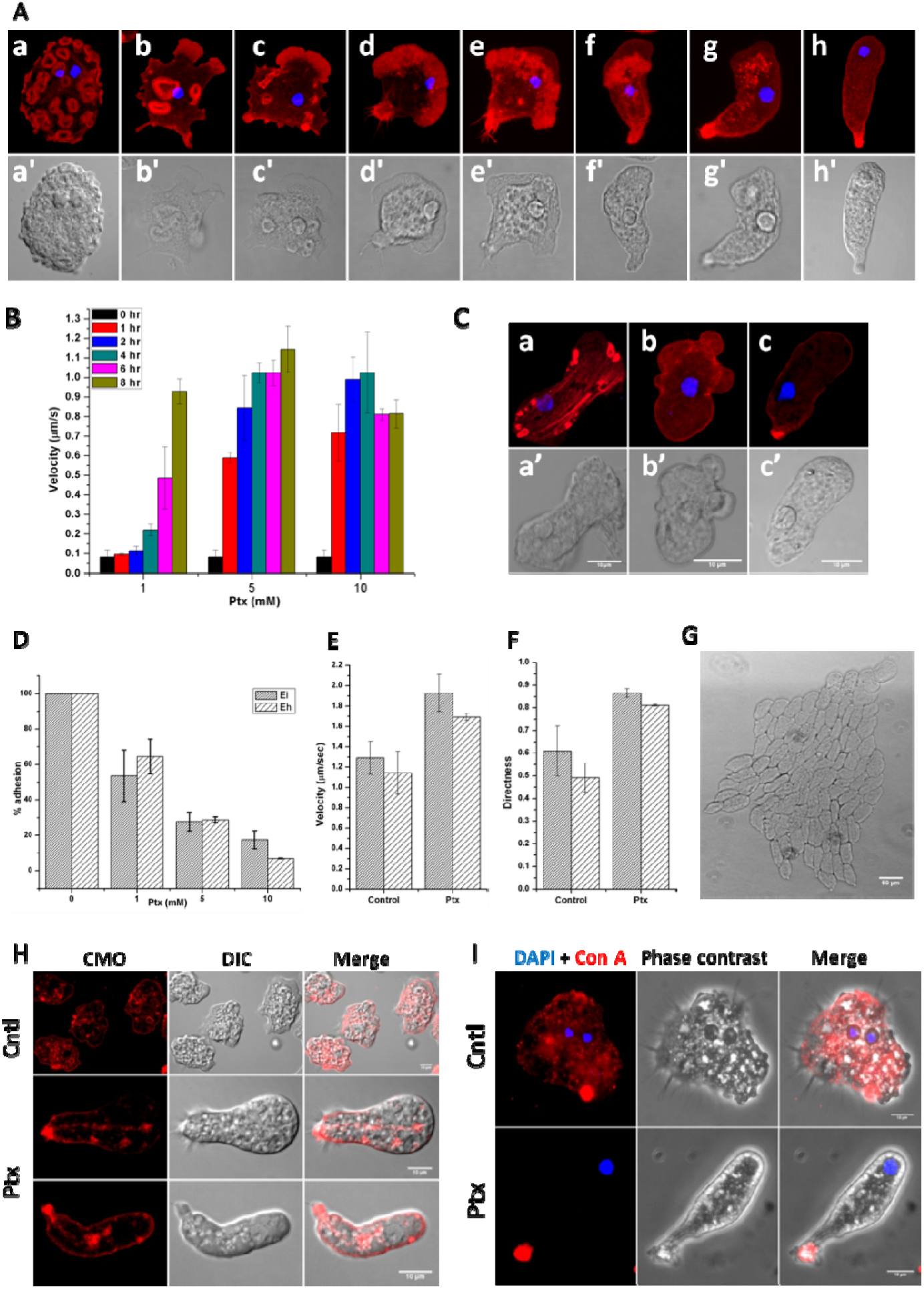
Morphological changes in response to pentoxifylline (Ptx). **(A)** In presence of 1 mM Ptx, adherent *E. invadens* trophozoites (a, a’) formed lamellipodial driven form (b, b’ to d, d’) followed by adherent bleb driven form (e, e’ to g, g’) and finally the non-adherent stable-bleb driven form (h, h’). (Upper panel: actin + nucleus, lower panel: DIC). **(B)** The change in cell speed with time after Ptx treatment. **(C)** In the presence of 10 mM Ptx, adherent *E. histolytica* trophozoites (a, a’) formed multiple blebs on the surface (b, b’) and then formed the stable bleb morphology (c, c’). **(D)** Ptx dose-dependently reduced cell adhesion. **(E)** Velocity and **(F)** Directness during motility under the agarose with and without Ptx. **(G)** The polarized cells moved collectively in a head to tail alignment under the agarose. Scale bar: 50 μm. **(H)** Plasma membrane staining of control and polarized cells with CellMask Orange (CMO). **(I)** Membrane proteins stained with fluorescent concanavalin A in control and polarized morphology. Scale bar: 10 μm.

Ptx was able to reduce cell adhesion dose-dependently (Fig. 3D), but these adhesion-deficient cells showed normal locomotion under confinement. During Under-agarose assay, the velocity of both *Eh* and *Ei* cells, under the agarose was found to be ~1 μm/s even in the absence of Ptx (Fig.3E Control). The multinucleated giant cells of *Ei* have shown a similar increase in motility under confinement (Krishnan and Ghosh, 2018) and this could be due to the induction of bleb-driven movement by mechanical resistance (Zatulovskiy *et al*., 2014). While velocity increased, motility remained random with directness values of 0.60 and 0.49 for *Ei* and *Eh*, respectively (Fig.3F, control). In the presence of Ptx, the velocity under the agarose was further increased to 1.9 μm/s in *Ei*, and to 1.69 μm/s in *Eh* (Fig.3E Ptx) and the motility became highly directional (D=0.8) for both the species (Fig.3F, Ptx). Under agarose, the polarized *Ei* were always found to move collectively after aligning themselves in head-to-tail fashion (Fig.3G, Movie 10) similar to the streaming in *Dictyostelium* (McCann *et al*., 2010). This indicates that *Ei* must be secreting some chemo-attractant, which attracted and aligned other cells in a head-to-tail fashion, but such alignment was not observed in *Eh*.

Since the stable bleb-driven morphology was adhesion deficient, we tried to find how these cells move. It has been shown previously that adhesion-independent amoeboid migration is driven by a rearward flow of plasma membrane (O’Neill *et al*., 2018). When the unpolarized cells were stained with plans membrane stain CellMask Orange (CMO), most of the fluorescence was observed on the membrane and internal vesicles (Fig. 3H, control). In the polarized form, fluorescence was mostly observed on the uroid (Fig. 3H, Ptx). Large numbers of fluorescent vesicles were observed originating from uroid and moved forward through the endoplasm, and these could be replenishing the anterior cell membrane. The membrane movement was also observed by the rearward flow of membrane proteins by treating the cells with fluorescent concanavalin A (ConA) (Fig. 3I). When unpolarized cells were briefly incubated with fluorescent ConA, most of the fluorescence was observed on the cell surface (Fig.3I, control). In the polarized morphology, the entire fluorescent ConA accumulated at uroid (Fig.3I, Ptx) which also indicated the rearward flow of membrane.

### Pentoxifylline effects are mediated by adenosine receptors

The biological effects of Ptx might be due to either its action on adenosine receptors or inhibition of phosphodiesterases. Like Ptx, adenosine was also found to be chemokinetic in Ei, since cell velocity increased to 0.50 μm/s (Fig.4A) when adenosine (0.1-1 mM) was added to the medium. The most important feature of adenosine-induced chemokinesis was the highly random movement as observed from the low directness value of 0.3 (Fig.4B) and the migration tracks (Fig.4C, Movie 11). In the presence of adenosine, protrusions were formed all over the cell surface in *Ei* (Fig.4D) causing the cells to move randomly in all directions, and Rhodamine phalloidin staining showed these protrusions were blebs (Fig.4E). These observations made in *Ei* were not clearly visible in *Eh* due to its isotropic blebbing and inherent random motility (Fig.4A, B). Thus the effects of adenosine are exactly opposite to those of Ptx, indicating both could be binding to the same *Entamoeba* adenosine receptor, which is not yet identified. Simultaneous stimulation of *Entamoeba* multiple adenosine receptors produced multiple leading edges causing the cell to move in opposing directions (Fig.4F), and so there was no productive cell motility. In contrast, Ptx treatment produced a single leading edge resulting in a polarized morphology causing the cell to move in a single direction (Fig.4G). These observations could be explained based on the mutual inhibition between frontness signals and backness signals (Xu *et al*., 2003; Wang *et al*., 2013). Adenosine seems to promote a strong frontness, whereas Ptx, a strong backness. Ptx and adenosine-induced motility patterns represented two extremes of *Entamoeba* morphology and the motility of trophozoites in growth medium over a long time shows elements of both randomnesses, and directional persistence (Fig.4H).

**Fig. 4.**
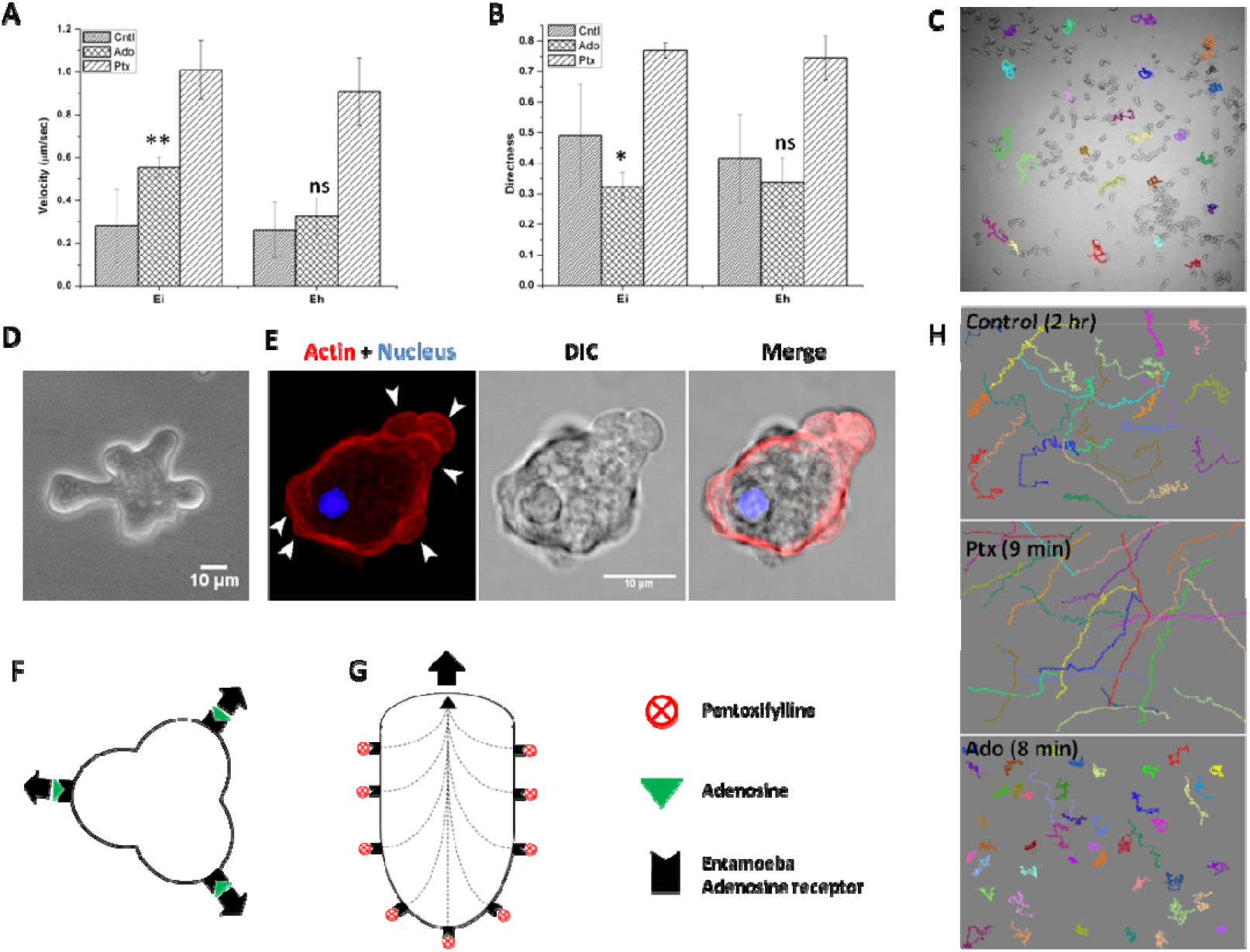
Adenosine induces opposite motility characteristics to pentoxifylline. Comparison of **(A)** Velocity and **(B)** Directness of control, adenosine treated and Ptx treated *E. invadens* (Ei) and *E. histolytica* (Eh). **(C)** The migratory tracks of adenosine treated cells. **(D)** Multiple protrusions on the adenosine treated cells. **(E)** Actin staining showing the presence of a large number of blebs (arrow head) all over the cells. **(F)** Adenosine binding to *Entamoeba* adenosine receptor activates frontness and produces multiple leading edges causing the cell to move in multiple directions. **(G)** Ptx binding activates the backness which then confines front to a single position and the cell moves in a single direction. **(H)** Comparison of motility using migratory tracks showed that the movement in growth media contained patterns of both pentoxifylline and adenosine-induced motility. Scale bar: 10 μm.

While *Ei* formed the polarized morphology in response to stresses and during encystation, in *Eh* such morphology was reported only under a chemoattractant gradient like that of TNF (Blazquez *et al*., 2006). When *Eh* and *Ei* trophozoites were exposed to continuous stress (oxidative or temperature), large numbers of polarized cells with a fluid-filled hyaline cap at the leading edge were observed (Fig.S4; Movie 12). These cells were similar to the stable bleb driven form, morphologically, and as per actin distribution pattern, but its importance is not yet understood. The polarization and motility of *Entamoeba* are required for invasive amoebiasis (Dufour *et al*., 2015; Aguilar-Rojas *et al*., 2016). Dominant positive expression of URE3-BP in *Eh* induced an elongated morphology which was reported to be highly invasive (Gilchrist *et al*., 2010), and so the polarized form may have an important role in the pathogenesis. The role of adenosine receptors on cell motility indicted the possibility of purinergic signaling (Chen *et al*., 2006; Ledderose *et al*., 2016) in controlling motility and chemotaxis of *Entamoeba*. Animal cells contain 5-10 mM of ATP in their cytosol, and the tissue damage by *Entamoeba* may cause leakage of ATP from damaged cells. This could lead to accumulation of purine nucleotides in the surroundings of *Entamoeba* and may influence the *Entamoeba* motility, chemotaxis and thus its pathogenicity. This is the first report of purinergic signaling in protozoan parasites and its possible involvement in invasiveness. In conclusion, *Ei* or *Eh* may be used as a model for studying different aspects of cell motility as its morphology can be easily controlled *in vitro*. Controlled motility and morphology will also be helpful to study the pathogenicity of amoebiasis.

## MATERIALS AND METHODS

### Cells and reagents

*Entamoeba invadens* (strain IP-1) and *Entamoeba histolytica* (strain HM1-IMSS) were cultured in TYI-S-33 medium completed with 10% heat-inactivated adult bovine serum (HiMedia) and 3% Diamond vitamin mix and penicillin and streptomycin at 25°C and 37°C, respectively. Pentoxifylline, caffeine, Texas Red-concanavalin A, DAPI, adenosine, and calcofluor white were purchased from Sigma-Aldrich. Rhodamine conjugated phalloidin, and CellMask Orange were purchased from Molecular Probes, Invitrogen, USA. Transwell inserts with 8μM pore were obtained from SPL biosciences.

### Cell staining

Clean coverslips were placed in 35 mm cell culture plates, and trophozoites at appropriate cell density were added and allowed to adhere. After the experiments, the cells were fixed using pre-warmed 3% paraformaldehyde for 30 minutes, followed by permeabilization with 0.2 % (v/v) Triton X-100 in PBS for 5 minutes. Nucleus was stained with DAPI, and chitin wall was stained with calcofluor white. The actin cytoskeleton was visualized by staining the cells with Rhodamine conjugated phalloidin after blocking the permeabilized cells with 2% (w/v) BSA. Texas Red-concanavalin A (50 μg/ml) and CellMask Orange (1:1000 dilution) were used to study membrane dynamics during motility.

### Microscopy

Time-lapse video microscopy was carried out using Olympus IX51 inverted light microscope with a camera attachment and photo-editing software (Image Pro Discovery). Olympus FV1000 confocal microscope with Fluoview software was used for fluorescence imaging. The raw images were processed using ImageJ.

### Encystation

To prepare the encystation medium (LG 47), TYI-S-33 medium without glucose (LG100) was prepared and diluted to 2.12 times and then completed with 5% adult bovine serum, 1.5 % vitamin mix and penicillin and streptomycin. To find the response of *E. invadens* to encystation media, cells were grown in tissue culture plates, and TYI-S-33 medium was replaced with the LG47 medium. *E. invadens* cultures below confluency and confluent cultures with small cell aggregates were used. The formation of chitin positive cysts was observed by staining with calcofluor white.

### Calculation of cell shape descriptors

The changes in the morphology were quantitatively defined using the shape descriptors like aspect ratio and circularity. From the images, ImageJ was used to calculate the shape descriptors, aspect ratio, and circularity. Aspect ratio is the ratio of the length of the major axis to the minor axis for this ellipse. For spherical morphology, the aspect ratio = 1, whereas for an elongated morphology the aspect ratio is >1. Circularity is calculated as 4π × [Area]/[Perimeter]^2^. Values of 1.0 indicate a perfect circle, and lower values indicate an increasing formation of protrusions. Stationary un-polarized cells showed high circularity, and low circularity is a trait of polarized migrating cells.

### Analysis of Cell Motility

Time-lapse video microscopy was performed using Olympus IX51 inverted microscopes with a camera attachment and photo-editing software (Image Pro Discovery). The images were then taken at appropriate intervals to produce a time-lapse movie. Using the ImageJ software with plugins like Manual Tracking and MTrackJ, the position of cells at different time points identified from time-lapse. An average of 20-30 cells was tracked during each experiment, and each experiment was repeated minimum three times. Then using the Chemotaxis and Migration Tool (Ibidi), the position of each cell was used to visualize the cell tracks, and from this, migration velocity and Directness were calculated. The velocity of each cell was calculated as the total distance covered by a cell divided by time, and the average velocity for each experiment was calculated. The Directness (D) is the measure of a cell’s tendency to travel in a straight line. It is calculated by dividing the displacement with the distance. D is the average of all values of directness for all cells. A directness of D = 1 indicates a straight-line migration from start to endpoint while D≪1 indicates random motility.

### Transwell migration assay

Transwell migration assays (Boyden Chamber Assay) were conducted to measure the chemokinesis and chemotaxis. For this Transwell inserts containing 8 μm pore polycarbonate membranes and 24 well-cell culture plates were used. To measure chemotaxis, complete TYI-S-33 medium or TYI-S-33 medium containing chemoattractant (0.5 ml) were placed in the bottom chamber. Trophozoites at a concentration of 1.0 x 10^5^ cells/ml in incomplete TYI medium were then added to the upper well chamber. To test chemokinesis, the chemical was separately added to the top and bottom chambers. The 24 well plates were then sealed with parafilm and placed in an anaerobic bag for 3 hours. Trophozoites migrated to the lower chamber were detached from the culture plate by keeping the plate on the ice, harvested, and counted using a heamocytometer. The extent of migration was expressed as the fold change of the number of migrated cells relative to control cells

### Under-Agarose assay

For Under-Agarose assay, 0.7% agarose dissolved in 100% LG was poured into 100-mm plastic Petri dishes and allowed to solidify. A 2-mm-wide trough was made at the center of the Petri dish. *Entamoeba* cells were harvested, numbers were adjusted to 10^5^ amoebae/ml in LG100 was added to the trough. Plates were maintained at 25°C and 37°C for *E. invadens* and *E. histolytica* respectively. Once the cells moved under the agarose gels, their movements were recorded with time-lapse microscopy, and the motility characteristics were analyzed as previously described.

### Statistical Analysis

Quantitative data are presented as the mean ± standard deviation of a minimum of three independent experiments. The significance of the experimental data was estimated using Student’s t-test, and the results were considered statistically significant only if p < 0.05. Statistical significance is represented as *p < 0.05, **p < 0.01, ***p < 0.001, ns not significant.

## Acknowledgments

DK is the recipient of Senior Research Fellowship from Indian Council of Medical Research, India. SKG acknowledge DBT, Govt. of India for partial funding. Authors thank FIST, DST, Govt. of India for the confocal facility.

## Author contributions

DK and SKG designed research; DK performed research; DK and SKG wrote the paper.

## Supplementary Figures

**Fig. S1.**
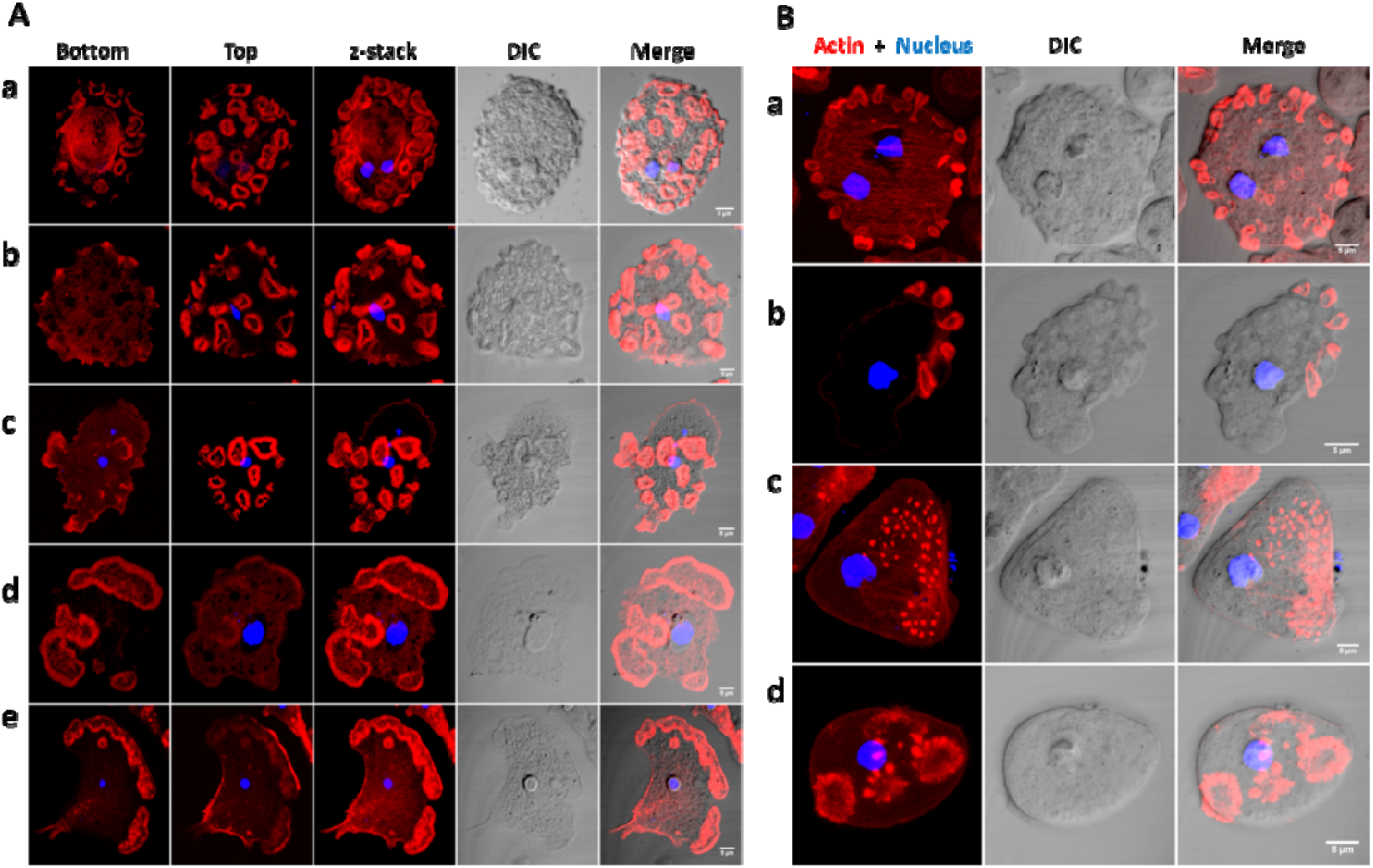
Cellular localization of F-actin in *E. invadens* and *E. histolytica* trophozoites. **(A)** In *E. invadens* trophozoites, actin is found as stress fibers at the ventral side of the cells and phagocytic and pinocytic invaginations (a, b, c). In some cases podosome rosettes were observed (d, e). **(B)** In *E. histolytica* it was found as stress fibers (a), phagocytic and pinocytic invaginations (b), podosomes (c) and podosome rosettes. Scale bar: 5 μm.

**Fig. S2.**
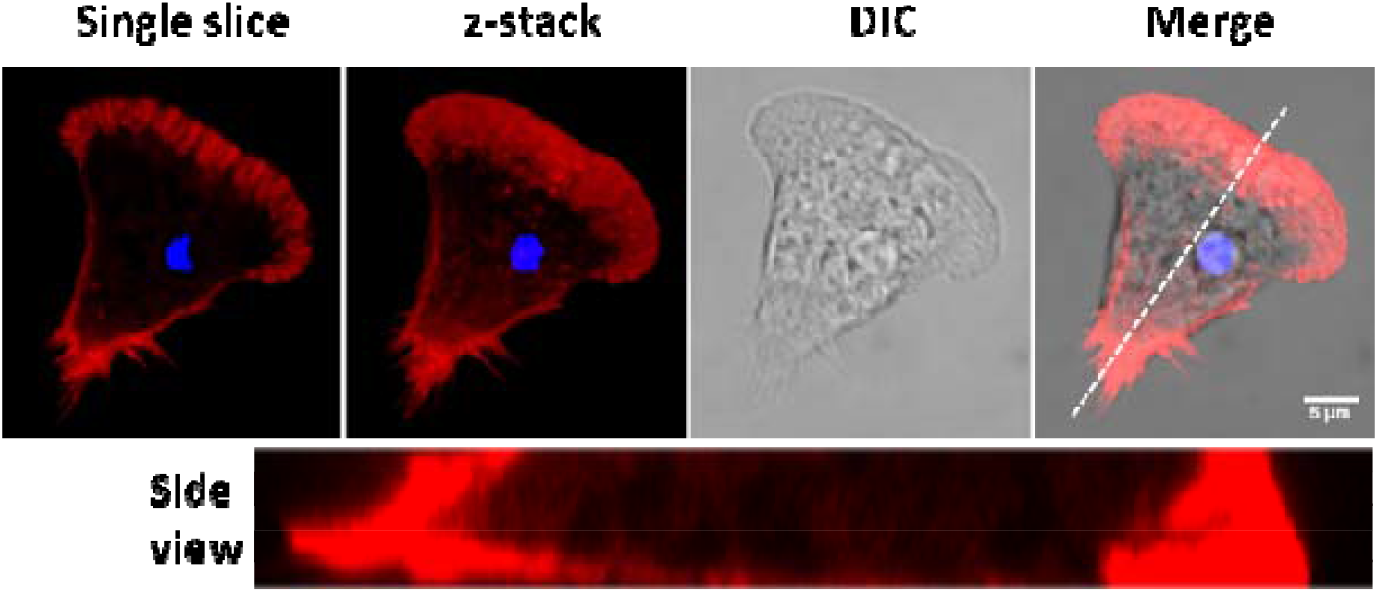
Lamellipodial motility. Lamellipodia based motility which is characterized by actin rich protrusions. The side view of cell is across the drawn line. Scale bar: 5 μm.

**Fig. S3.**
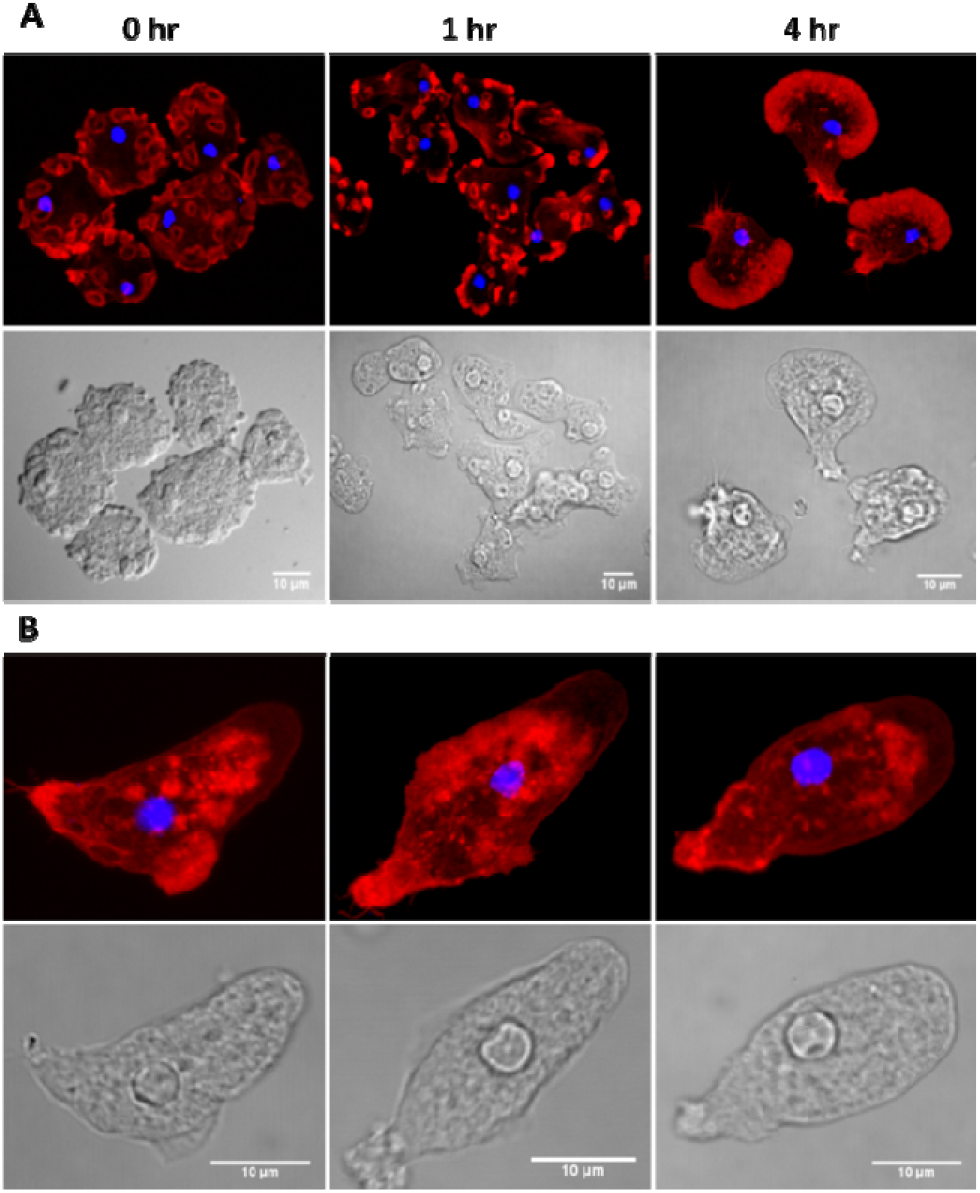
Intermediate morphological stages during the amoeboid transition identified by changes in F-actin localization. **(A)** In the un-polarized cells actin is found in the phagocytic and pinocytic invaginations. Within 1 hour of pentoxifylline treatment (1 mM) lamellipodia started appearing and the phagocytic invaginations started disappearing. Lamellipodia became very prominent by 4 hours and no phagocytic invaginations were observed. **(B)** By 4-6 hours as the cell became elongated, the lamellipodia disappeared and blebs appeared. These cells had large number of actin dots on their ventral surface which could be adhesion structures. Scale bar: 10 μm.

**Fig. S4.**
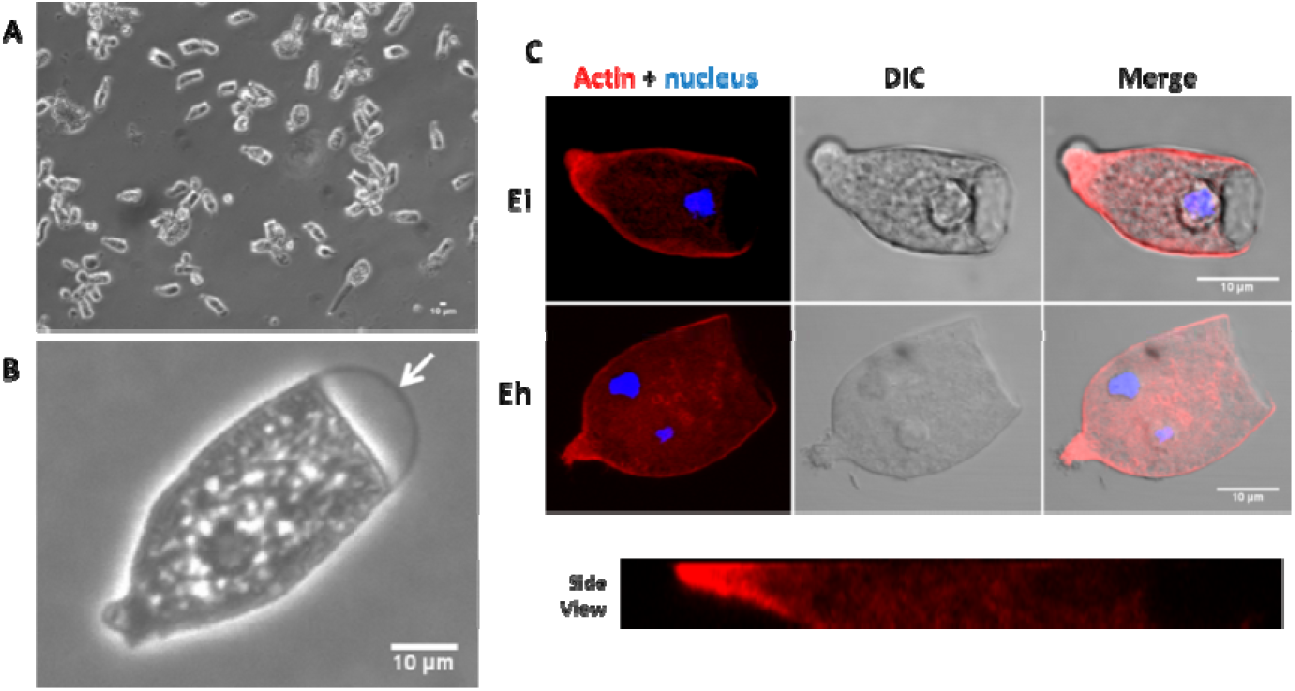
Polarized morphology fluid filled hyaline cap. **(A, B)** When exposed to continuous stress *Entamoeba* formed an amoeboid form with fluid filled hyaline cap at the leading edge containing no actin **(C).** Scale bar: 10 μm.

## Supplementary Movies

**Supplementary Movie 1.**
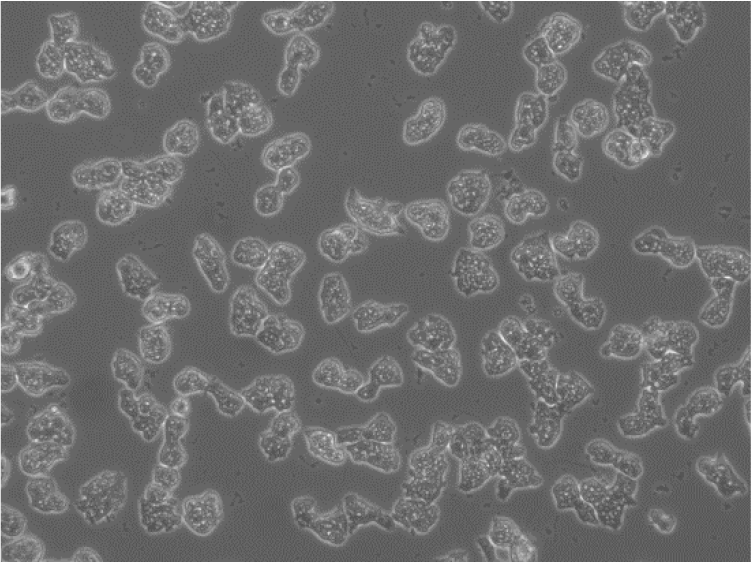
Morphological and motility changes induced by Pentoxifylline.

**Supplementary Movie 2.**
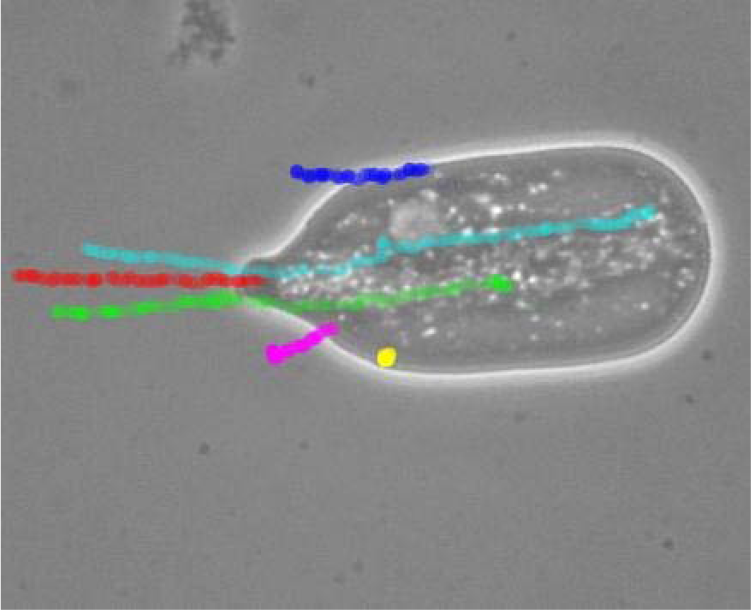
The cytoplasm of polarized form was divided into a fluid endoplasm and a gelled ectoplasm as shown by the speed of particles (coloured tracks) compared to the cell (red).

**Supplementary Movie 3.**
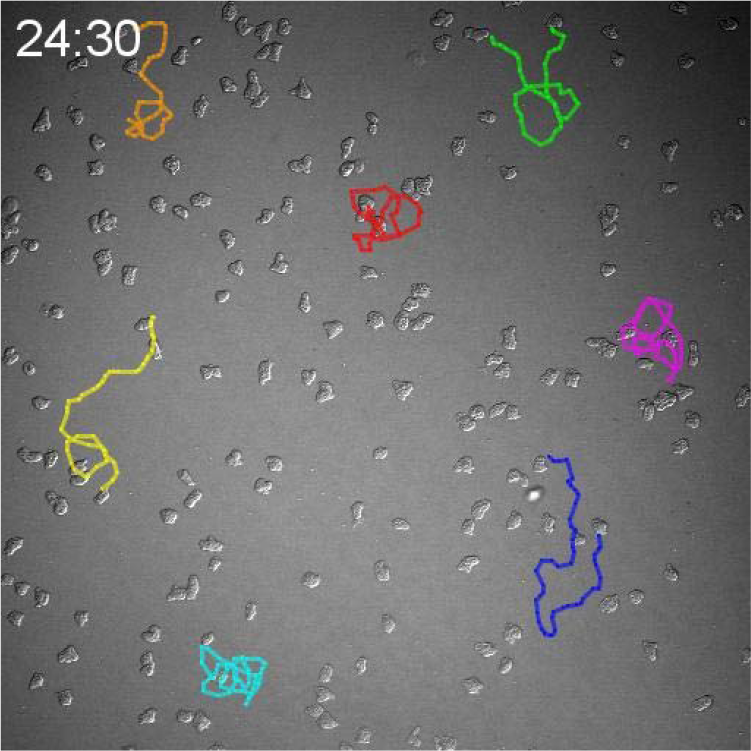
Motility pattern of unpolarized and polarized cells of *E. invadens* shown by cell tracks.

**Supplementary Movie 4.**
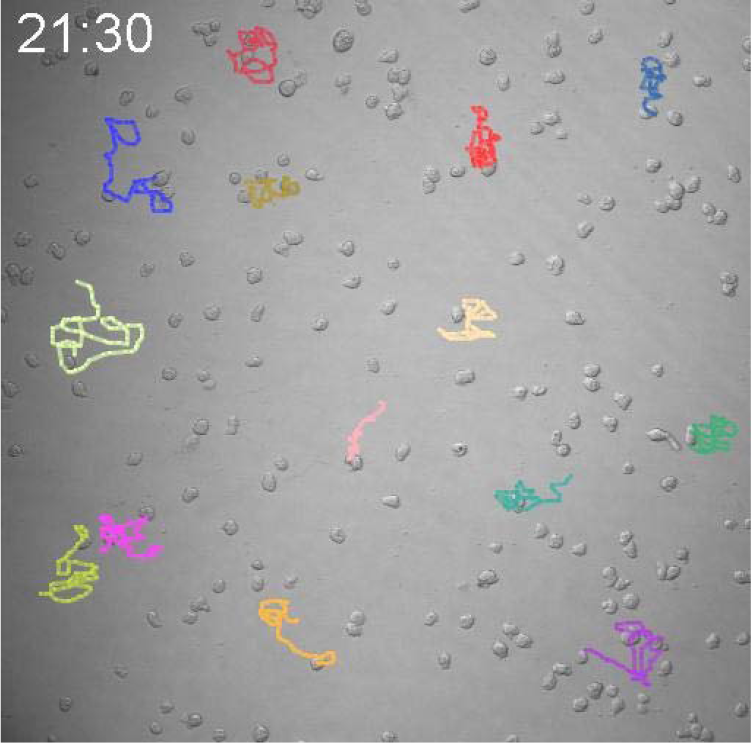
Motility pattern of unpolarized and polarized cells of *E. histolytica* shown by cell tracks.

**Supplementary Movie 5.**
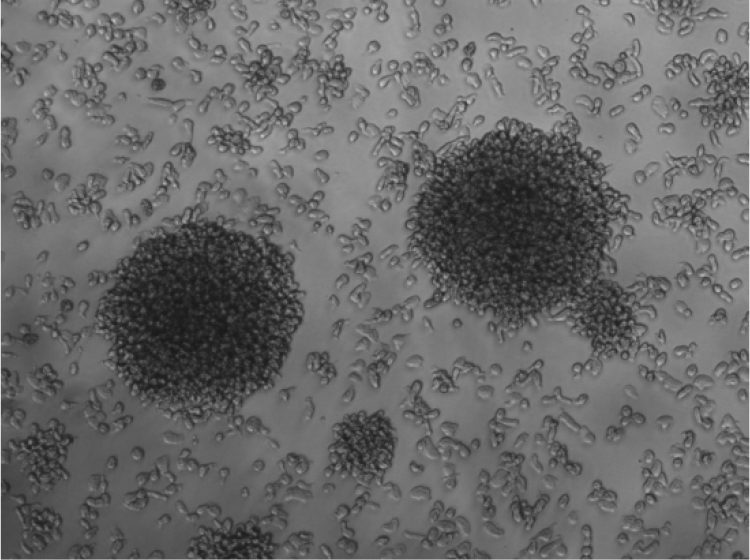
The pentoxifylline induced cell aggregates was galactose ligand mediated as they were dispersed by 10 mM Galactose.

**Supplementary Movie 6.**
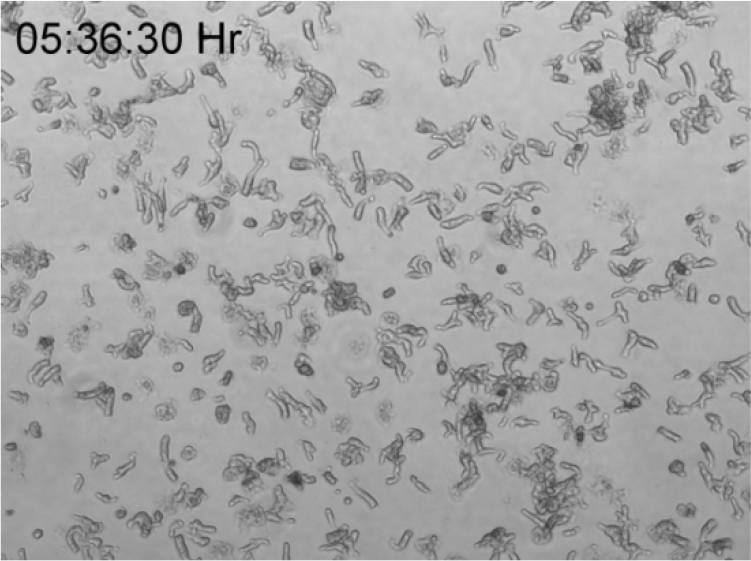
Cell polarization in response to encystation media.

**Supplementary Movie 7.**
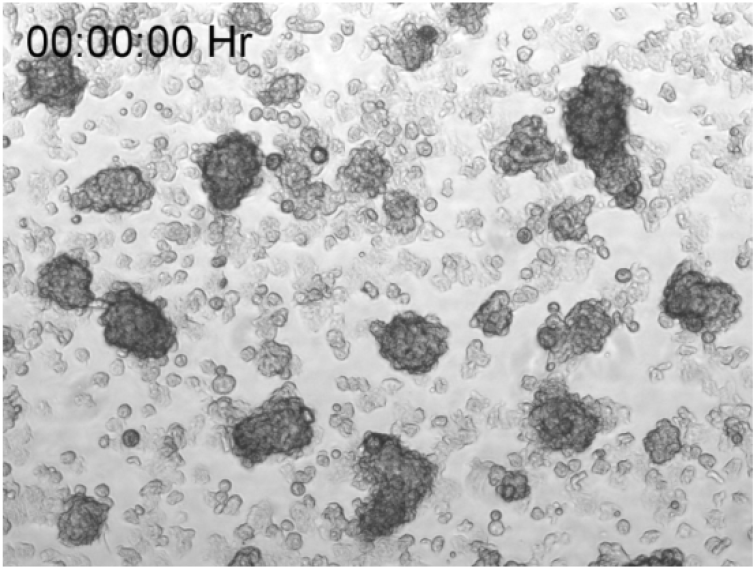
Migration of polarized morphology towards smaller aggregates response to encystation media.

**Supplementary Movie 8.**
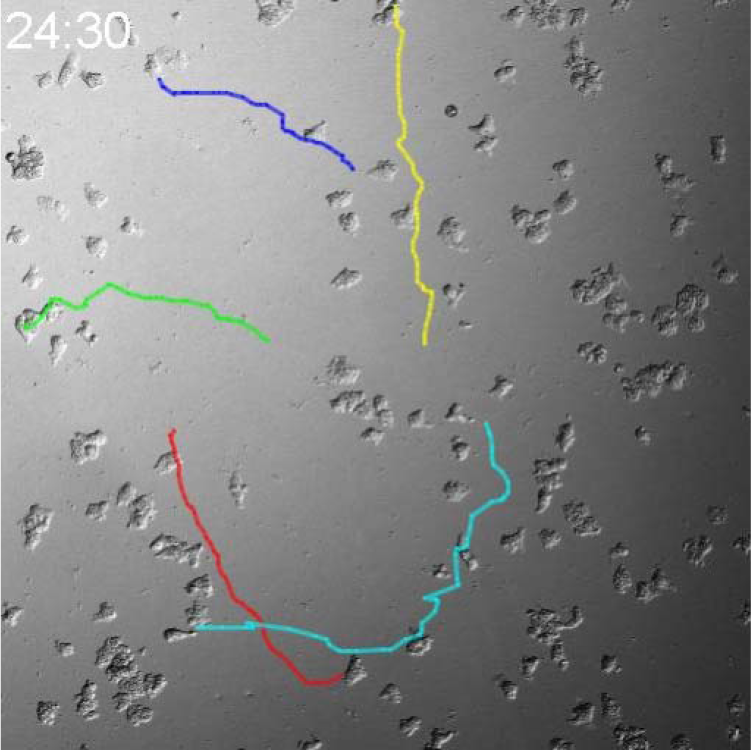
Lamellipodial motility during early stages of cell polarization.

**Supplementary Movie 9.**
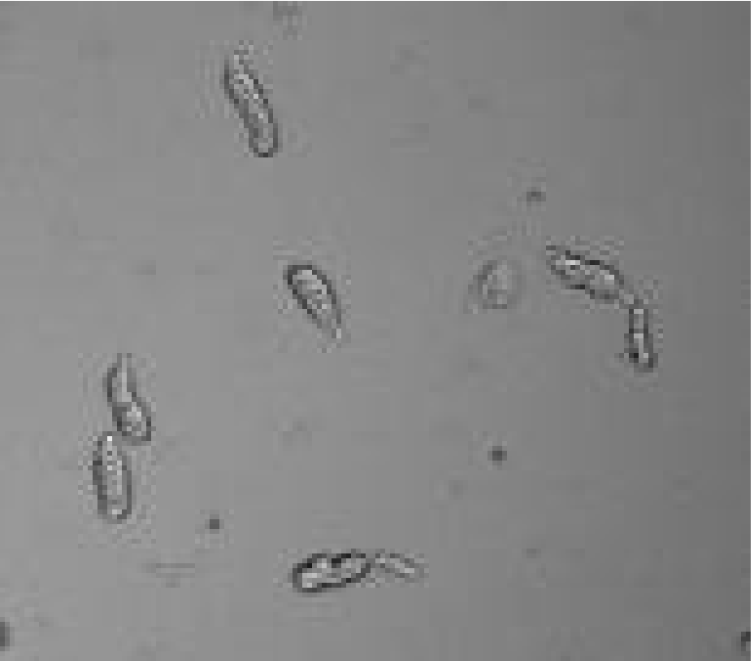
The polarized morphology did not show any cell adhesion causing it to slip during movement.

**Supplementary Movie 10.**
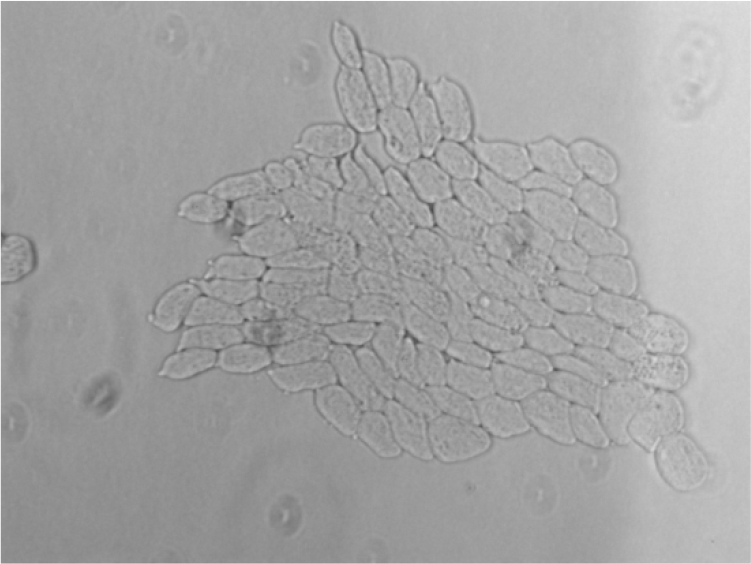
The polarized cells moved collectively in a head to tail alignment under the agarose.

**Supplementary Movie 11.**
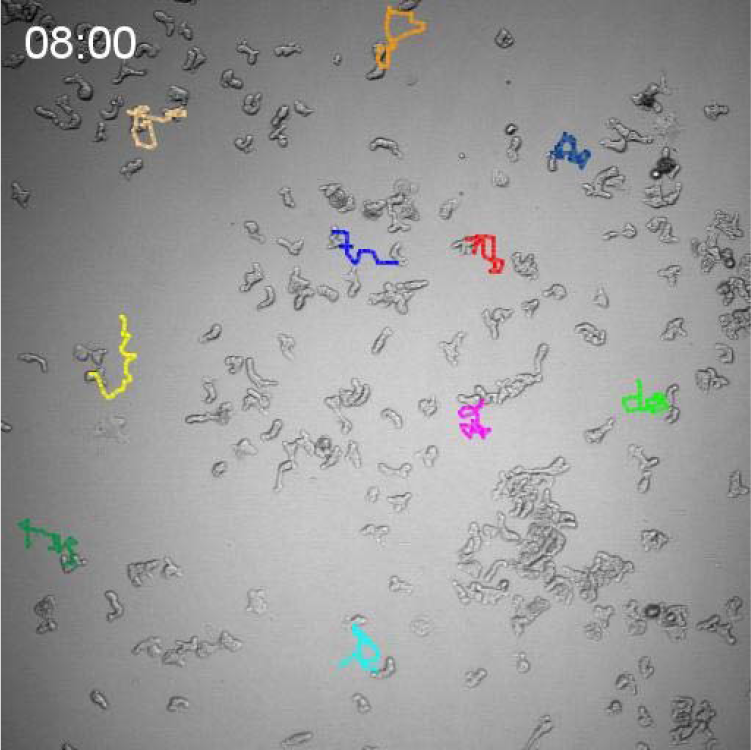
Motility pattern of adenosine treated cells of *E. invadens* shown by cell tracks.

**Supplementary Movie 12.**
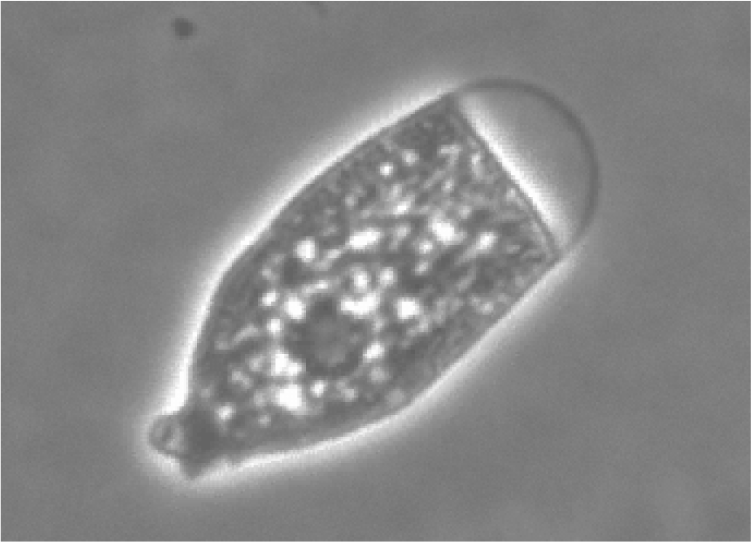
When exposed to continuous stress *Entamoeba* formed an amoeboid form with fluid filled hyaline cap at the leading edge

